# An Open-Hardware sample mounting solution for inverted light-sheet microscopes with large detection objective lenses

**DOI:** 10.1101/636977

**Authors:** Craig T. Russell, Eric J. Rees

**Affiliations:** Dept. of Chemical Engineering and Biotechnology, Cambridge University, Cambridge, U.K.; The National Physical Laboratory, London, U.K.

**Keywords:** Light-sheet microscopy, Open-Hardware, Sample mounting, inverted light-sheet microscopy, 3D printing

## Abstract

Implementations of light-sheet microscopes are often incompatible with standard methods of sample mounting. Light-sheet microscopy uses orthogonal illumination and detection to create a thin sheet of light which does not illuminate the sample outside of the depth of field of the detection axis. Typically, this configuration involves a pair of orthogonal objectives which constrains the positioning of a length of coverslips or microscopes in range of the detection objective. We present an open-hardware (1, 2) sample mounting system for light-sheet microscopes using large detection objectives.

---

During the past decade light-sheet technology has emerged as alternative to the well-established confocal microscope. A significant drawback of Light-sheet Fluorescence Microscopy (LSFM)s is that sample mounting can be cumbersome. There have been several attempts at imaging on flat glass in order for light-sheet microscopes to be compatible with favoured sample mounting techniques. The simplest method involves using a pair of objectives with an aperture angle of less than 90° and mounting those above a sample holder. Fig. 2a shows how the di-SPIM (3) uses a pair of 40 × 0.8 NA objectives to allow a glass slip to be inserted, whilst sacrificing NA when compared to larger detection objective lenses. The NA is then reclaimed through image fusion, albeit after image data for two volumes are acquired from two cameras at a cost of halving temporal resolution. The lattice light sheet(4), for instance, chooses a special objective pair such that the sum of its solid angles, seen in Fig. 1, of its numerical apertures does not interfere with the glass raised from below. To do so, a custom excitation objective is mounted along a with Nikon 1.1 NA 25× objective at angles to the sample mounting section that allows a sample chamber to be inserted, as shown in Fig. 2b.

**Fig. 1.**
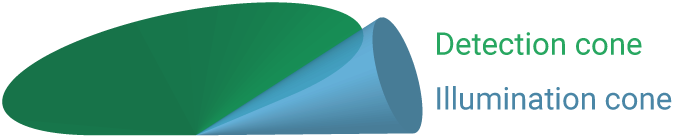
The combined solid angles of two objectives must not exceed 180° in the plane of orthogonality of the objectives.

**Fig. 2.**
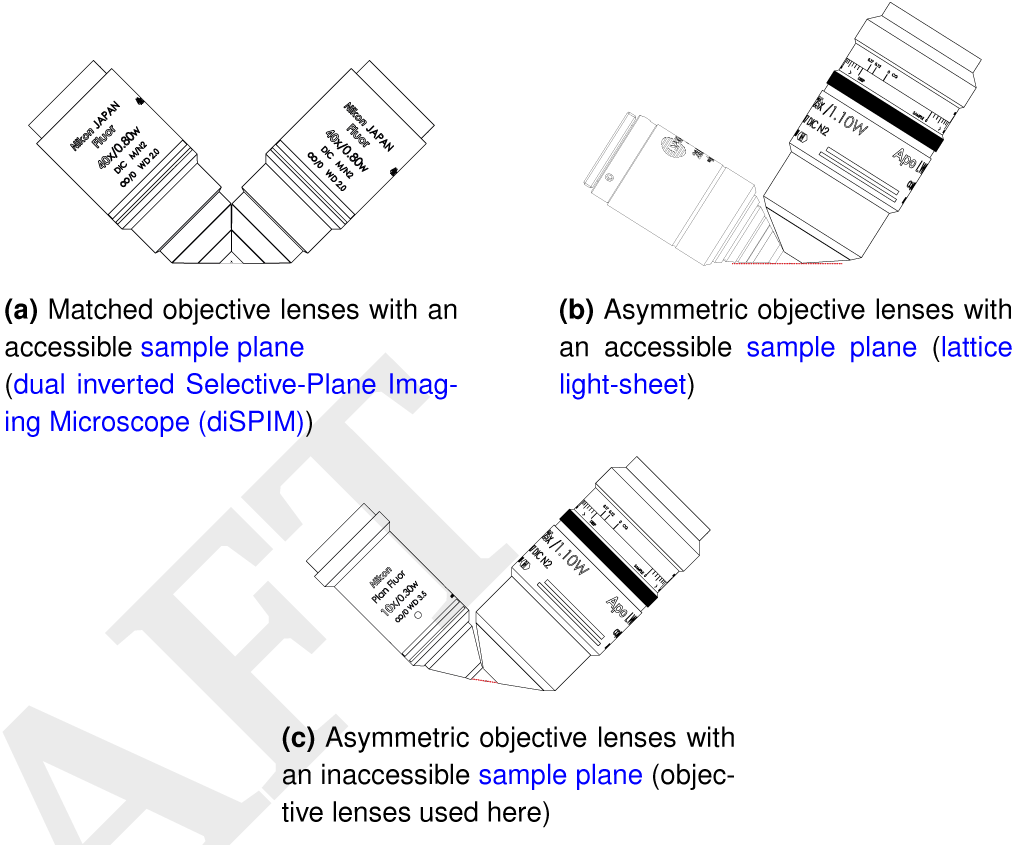
Sets of objective lens pairs for large NA light-sheet microscopy, dotted lines show available sample planes.red from two ca (a) shows the objective arrangement as found in the diSPIM, whereby each imaging arm is computationally fused to create a final image volume with higher axial resolution, from the two views. (b) shows the objective arrangement as found in the lattice light-sheet, the objective lenses are chosen to maximise the imaging NA and the illumination NA whilst allowing the sample plane to be accessible (c) shows the objective combination used in this work, where the imaging and illumination objectives both have a high NA, however the sample plane is inaccessible

As depicted in Fig. 4 and Fig. 3, other techniques for light-sheet imaging require specialist sample mounting and preparation to be employed. These methods present a barrier that can discourage new users of light-sheet microscopy; this paper presents protocols and open-source designs, using 3D printing technology, to help dispel this barrier.

**Fig. 3.**
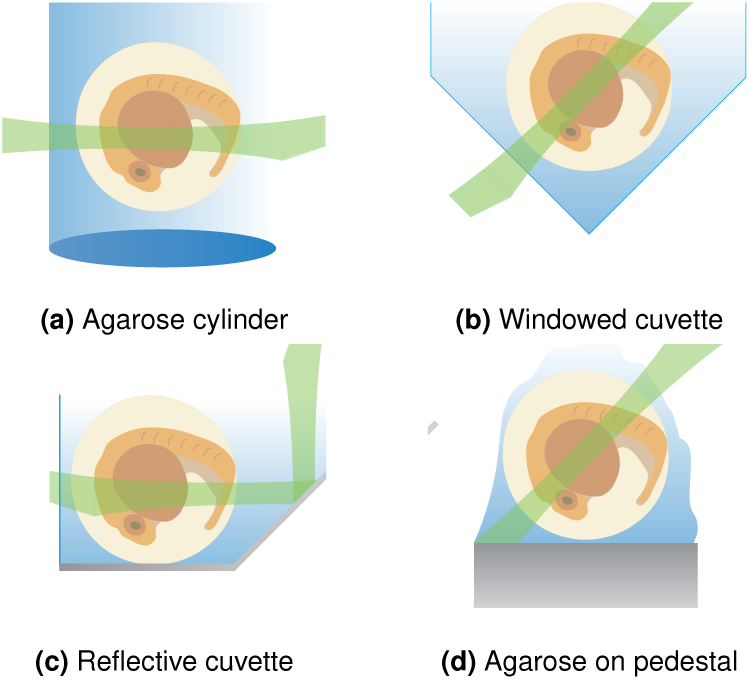
Strategies for mounting zebrafish are shown, though the techniques apply widely to other model organisms. (a) uses a cylinder of agarose suspended from above and lowered into the imaging volume; (b) is a cuvette whereby the flat windows are imaged and excited through orthogonally; (c) shows a reflective cuvette as in Fig. 4 (d) has a small pedestal with a small amount of agarose placed on top, the entire system is submerged in medium to match refractive indices.

**Fig. 4.**
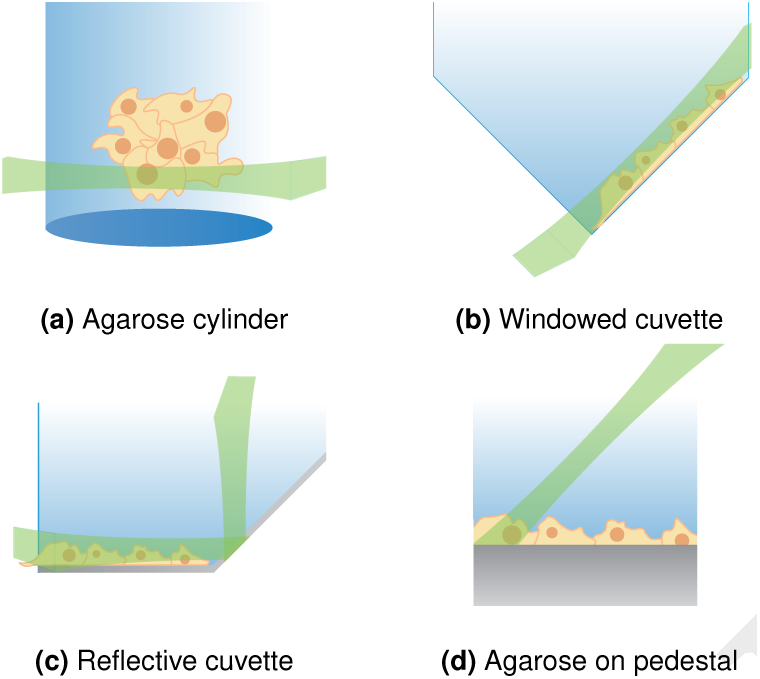
Strategies for mounting cell culture. (a) shows a cylinder of a gel used to scaffold a 3D cell culture. (b) Shows cells cultured on a cuvette with the illumination arriving through the orthogonal flat window. (c) shows a reflective cuvette being used to redirect a concomitant illumination and detection into orthogonal, similar reflection strategies exist such as mounting small mirrors to a single objective or positioning small mirrors carefully near the sample. (d) shows cells cultured on a small pedestal and imaged using an iSPIM configuration, difficulties arise in keeping such pedestals sterile during incubation and then attaching to the system whilst submerged.

3D printing allows the home user to print structurally integral and disposal shapes with the ability to reconfigure the shape to their needs. Here we present open source CAD files for a design that we have optimised for a sample chamber that is compatible with an inverted Selective-Plane Imaging Microscope (iSPIM). Our design allows for the imaging of cell culture and embryos in their native conditions for long term microscopy (1).

## The design and construction

The sample chamber presented here is designed around a pair of objective lenses, a 1.1 NA 25 × Long Working Distance (LWD) and a 0.3 *NA* LWD 10 × water-dipping Nikon objective (5). Both are used for physiological imaging, and are widely used in the field of light-sheet microscopy. Fig. 2c shows how this objective pair, if mounted at 45° to a flat sample will intrude on the imaging volume. Similarly the sample plane of the detection objective cannot be reached at any angle for the pair of objectives.

At the core of the design is a right-angle shelf on which a cover-glass bearing the specimen is mounted on This gives access to 2 mm× 20 mm area of accessible cover-glass at of one of its edges. Beneath the shelf is a recess which allows for sample positioning from below using a Raspberry Pi-Cam, as well as mitigating any reflection or scattering from the laser illumination source, Fig. 5a presents an an illustration of the recess. The sample is held down firmly by a 3D printed brace with embedded magnets. The brace is coupled to magnets inserted below mounted in a printed tray, which inserts into a slot. The slot also allows for heating module, making it suitable for cell culture, embryos, microbes and more. The temperatures used will not damage the printed chamber material (Acrylonitrile Butadiene Styrene (ABS) plastic printed using a MakerBot Replicator 2x) begins to deform at 140°.

**Fig. 5.**
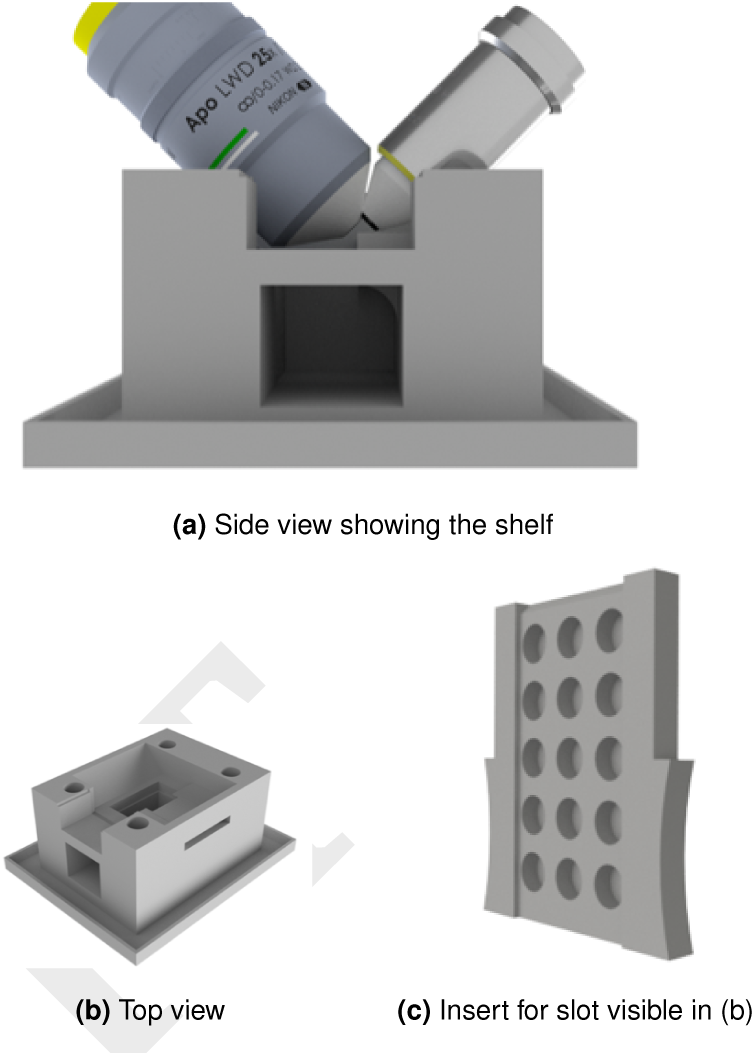
Computer Aided Design (CAD) model of proposed sample chamber, featuring modular heater inserts with magnet cut-out tray. On the side of the chamber and below the sample, Perspex windows are inserted and sealed using cyanoacrylate (Super-Glue), this ensures a liquid-tight seal. (a) shows the void below the sample mounting shelf for the camera as well as the view used by the user to position the sample. (b) is the top view showing the chamfered edges used for medium minimisation and as physical limits to protect the objectives. (b) also shows the 4 mounting through-holes for fixing the chamber to a stage and the drip stray below. (c) is the heater and magnetic mounting insert, small neodymium magnets insert into the holes and a heater pad is placed on top.

Perspex windows are laser cut and inserted in the side of the chamber (for user viewing) and below the sample (for additional illumination and camera viewing). Cyanoacrylate (Super-Glue) is used to secure the windows to the sample chamber, which ensure a liquid-tight seal. The adhesive is applied whilst the surface’s protective film is attached to the Perspex and the film is removed only when the cyanoacrylate is fully set, as cyanoacrylate vapour settles on surfaces and causes opacity. This allows both objectives to be accurately positioned, with respect to the mounted sample, by eye, fine correction can then be achieved using texture in the sample either from bright-field or fluorescence modes.

The transparent sample window below not only allows for a small camera module (Pi-cam, Raspberry Pi (6)) to be positioned but also for the attachment of a supplemental light source (Bright-Pi (7)) which offers sufficient bright-field illumination as well as an infrared modality. The under-sample illumination is useful for the positioning of samples without laser illumination, minimising overall photon dose to the sample. Users may also find the additional illumination modes valuable when coarse positioning using a camera on the low magnification illumination arm. (In the present system, the illumination arm is a 10 × objective versus the detection magnification of 25 × 1.25 ×, and so coarse positioning can be performed using the illumination objective.)

The camera model chosen here, uses a very short focal length lens (4 cm), which is matched to the height of void below the sample-mounting in the sample chamber. It is also compatible with the Bright-Pi, which offers sufficiently bright field illumination as well as an infrared modality. These features could be used in conjunction with the appropriate userdefined correction filters in the detection paths. To protect the delicate objectives from touching the sharp edge of the coverglass, the sample chamber is chamfered along it edges parallel to the objectives, restricting their lateral movement. The chamfers also serve to minimise immersion liquid within the sample chamber, decreasing the cost per experiment as well as mitigating deleterious spillage. At the bottom of the sample chamber, an additional drip tray is incorporated to help prevent immersion medium spillage due to overfilling.

For cellular imaging, the entire assembly may be cleaned by immersing in Isopropanol or Virkon for several minutes; this can be supplemented with ultra-violet light sterilisation if required.

Chambers were printed from ABS and Polylactic acid (PLA) which are ubiquitous, and biologically inert, plastics used in 3D printing. It should be noted that ABS and PLA 3D printed chambers do not survive autoclaving, nor overnight washes with Vircon or IPA. If the chamber is exposed to severe imaging conditions (contagions, prions, toxic chemicals) that require containment, multiple chambers may be printed and disposed of; allowing for high-throughput imaging when compared to a machined chamber which would require sterilisation after each use. The design features cleared M6 holes to mount the chamber to a breadboard below. In the present design the chamber is attached to a breadboard insert on an XYZ translation stage. Other spacings and sizings of mounting holes may be easily added to the chamber ad-hoc, using standard Three Dimensional (3D) CAD packages (ex. Autodesk Inventor, SolidWorks, OpenScad).

## Sample preparation

In Fig. 6 zebrafish were manually dechorionated, with tweezers, on a bed of solid agarose immersed in embryo medium, using a seperate stereo-microscope for dissection. The dechorionated zebrafish were then transferred into molten agarose and gently drawn into a length of Fluorinated Ethylene Propylene (FEP) tubing attached to a pipette tip. Organisms are mounted in agarose within in FEP tubing to help match refractive indices between immersion medium and the agarose. 0. 8 % agarose VII was used as higher percentage agarose may restrict the growth of a developing embryo as well as cause more scattering and refractive index mismatch; agarose VII produces the best imaging conditions of the agarose (8). Once the embryo was embedded in its agarose tubing it was transported (made safe and convenient in the FEP) for imaging. It was then adhered to a 25 mm coverslip using colourless nail varnish at each end of the tubing as in Fig. 7b, whilst avoiding the active imaging area along the FEP tubing.

**Fig. 6.**
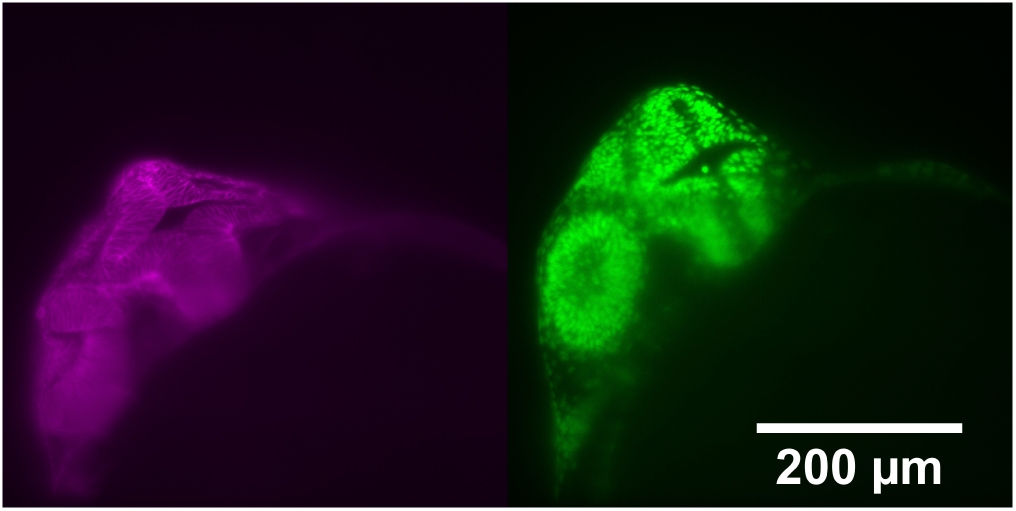
Two colour imaging of a transgenic zebrafish 24 h.p.f. The left image shows a membrane-local Beta-actin: mcherryCAAX probe; the right image shows fluorescent histones within the nucleus using a h2b:GFP probe.

**Fig. 7.**
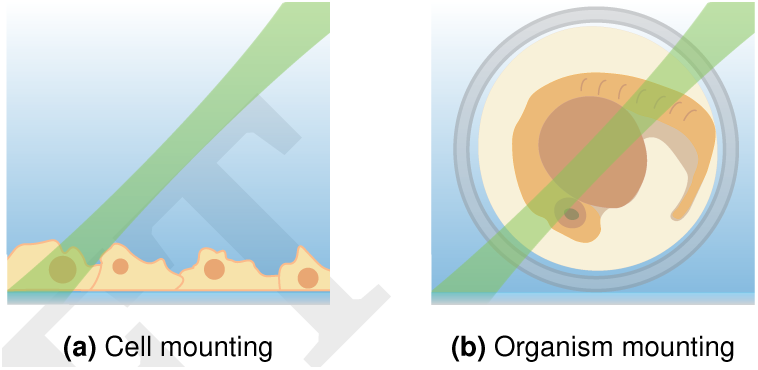
Strategies for mounting using the edge of a coverslip as proposed in this work. In (a), cells are cultured on cover glass for which protocols for sterile cell mounting are widespread. In (b), organisms are mounted in FEP tubing containing agarose and are adhered to cover glass using nail varnish.

For cell culture imaging, SH-SY5Y neuronal cells were cultured of on coverglass, as in Fig. 7a that had been functionalised with poly-l-lysene (which is positively charged and glass adherent) to encourage cell adhesion. In Fig. 8 cells were fixed using Formaldehyde and imaged in Phosphate-Buffered Saline (PBS). For live cell studies a medium without phenol red (fluorescent), and that does not require atmospheric control, should be used (ex. HEPES).

**Fig. 8.**
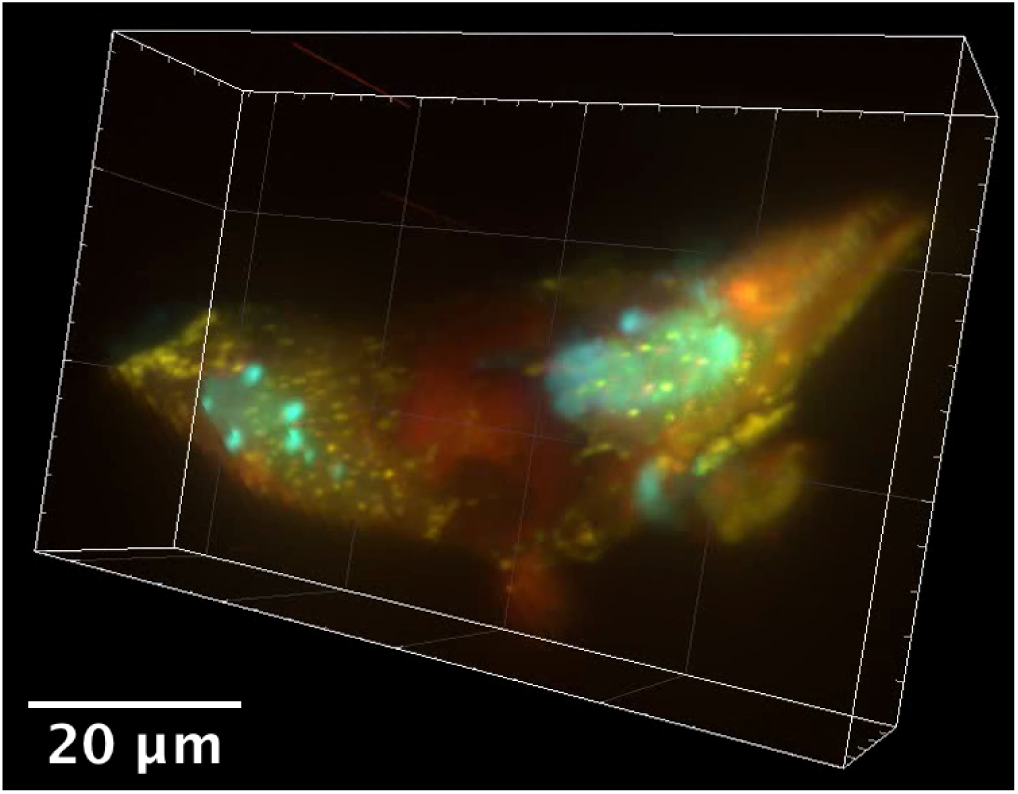
A three-colour composite 3D image, as rendered in Imaris, of a pair of SHSY5Y cells. The fluorescent stains used were m-Turqoise (blue), cYFP (yellow) and td-Tomato (red)

### Sample mounting and positioning

Samples mounted on coverslips are held in the chamber using the magnetically positioned bar to grip the coverslip. Pre-warmed immersion medium is added slowly and filled to 3 mm below the top of the chamber. The entire chamber is then driven below the objective pair by eye and carefully raised. Lateral positioning is best achieved by matching eye-level with the sample and adjusting the stage as required. Axial positioning is best coarsely adjusted by driving the sample slightly away from the laser illumination so as to not harm the sample. The scattered spot on the immersion-medium-glass interface should be minimised by eye. Finally, the sample chamber should be moved laterally, again, until a fluorescence signal on the camera can be detected. The secondary Raspberry Pi-cam below may also be used as an additional positioning camera. This is more useful for embryos, as the coarse adjustment method can be challenging due to early embryos being mostly transparent, even when using guiding marks on the coverglass. When mounting very large samples or samples that cover large areas, multiple images may need to be stitched together creating a mosaic. For the objective pair as used here, the sample chamber allows for 2 mm by 10 mm lateral and 1 mm axial movement. For systems with piezo objective scanners, the mosaicking of volumes may drastically reduce the overall time acquisition of a mosaicked volume by requiring fewer overall steps. Both approaches are available for the design presented here, provided care is taken to set hard limits on where the sample chamber is driven to by the automated stage.

## Conclusions

We have presented an open hardware solution to address the challenge of mounting a breadth of biological samples in large NA inverted light-sheet systems. The principles behind the design can be used directly, applied and extended to provide a robust starting point for which users can easily modify for their needs. Using a single piece of 3D printed material means the unit is robust (as it has no moving parts), disposable, sterile and mass producible. The unique design also allows for a large volume of travel, allowing for volumetric mosaicking and medium-throughput imaging. The device includes many useful features such as a sample mounting camera below, a Perspex window for precision positioning, multiple safety features and sample heating. Future models of this design could also include atmospheric control for long time-lapse imaging of cell cultures. Such a sample chamber may also be used for other biological samples at the same scale of cells to embryos

## Supporting information

Chamber design files

## ACKNOWLEDGEMENTS

C. Russell would like to acknowledge the Integrated Photonic and Electronic CDT for funding this work.

